# Simultaneous stabilizing feedback control of linear and angular momentum in human walking

**DOI:** 10.1101/2025.01.09.632072

**Authors:** Jaap H. van Dieën, Sjoerd Bruijn, Koen K. Lemaire, Dinant A. Kistemaker

## Abstract

Stabilizing bipedal gait is mechanically challenging. To analyze how gait is stabilized, previous studies have focused on the control of the body center of mass (CoM). These studies often linked deviations in linear momentum of the CoM to subsequent shifts in position of the center of pressure (CoP), or of the foot, relative to the COM, and interpreted these as stabilizing responses to correct linear CoM momentum. Mechanically, however, CoP shifts do not cause changes of linear CoM momentum, whereas they do cause changes in whole-body angular momentum. We hypothesized that experimentally observed correlations between CoP to CoM distance and horizontal ground reaction forces are related to the need to control both linear and whole-body angular momentum. We show that, in human walking, linear and angular momentum follow quasi-periodic functions with similar periodicity and phase. Combining the equations of linear and rotational motion for a system of linked rigid segments shows that, in this case, the horizontal distance between CoP and CoM should be correlated to horizontal force in the corresponding direction. This suggests that linear and angular momentum are simultaneously controlled and may explain the success of preceding studies that correlated CoM states to CoP or foot locations. Regression models fitted to experimental data of participants walking at normal and slow speeds showed that deviations in horizontal ground reaction forces and in moments of the ground reaction force about the sagittal and transverse axes could be predicted from deviations in the preceding linear and angular momentum respectively. Our analyses support that linear and angular momentum are indeed controlled simultaneously in human walking.

## Introduction

Stabilizing bipedal gait is challenging as illustrated by the high incidence of falls in toddlers learning to walk (Ghassabian et al., 2016) and in older and diseased adults (Robinovitch et al., 2013). The mechanical challenge of stabilizing bipedal gait arises from the body being upright and segmented and the small base of support. Additionally, the anatomy is “top-heavy,” with heavier segments supported by lighter segments. Consequently, perturbations, whether internal or external, are amplified by gravity, potentially causing instability. Robust gait, therefore, necessitates the control of the entire body through muscles delivering moments around the joints they span.

Conditions for stabilization of gait can be obtained from reduced order models, focusing on the “external coordination”, such as the control of the position or linear momentum of the body center of mass (CoM) relative to the base of support (De Comite and Seethapathi, 2024; Seethapathi and Srinivasan, 2019; van Leeuwen et al., 2022; Vlutters et al., 2016; Wang and Srinivasan, 2014) or of the angular momentum of the body around the CoM (Herr and Popovic, 2008; Silverman et al., 2012). Given that body mass can be considered constant, linear momentum and velocity are exchangeable; here we use momentum given the analogy between linear and rotational motion. Total forces and moments on the body determine the rates of change in linear and angular momentum and thus these variables will reflect all actions to stabilize linear and angular momentum respectively.

Hof (Hof, 2007) proposed mechanisms for the control of the linear momentum of the CoM based on the equation of rotational motion for the whole body modeled as a system of linked rigid segments. He rewrote the equation describing rotational motion around the projection of the CoM on the support surface CoM’ to:

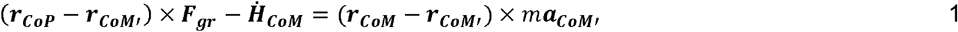

For conciseness, we here ignore forces other than gravity and the ground reaction force. Hof stated that the first term in equation 1 reflects the ‘move the center of pressure (CoP)’-mechanism. In similar vein, two of the authors of this paper (van Dieën and Lemaire, 2023) stated that this shows that shifting CoP relative to CoM’ can control linear momentum. This notion was derived from inverted pendulum models as well (Morasso et al., 1999; Townsend, 1985; Winter, 1995) and is reflected in many studies on stabilization of bipedal gait from our and other groups, whether explicitly using the position of the CoP, or the position of the foot relative to the CoM (De Comite and Seethapathi, 2024; Hof et al., 2010; Seethapathi and Srinivasan, 2019; van den Boogaart et al., 2022; van Leeuwen et al., 2022; van Leeuwen et al., 2021; Vlutters et al., 2016; Wang and Srinivasan, 2014). However, shifting the CoP does not affect linear momentum, as the equation of linear motion of a system of rigid linked segments is:

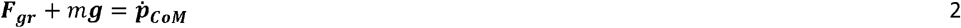

Evidently, the CoP does not appear in equation 2. Using equation 1, a shift in CoP predicts CoM acceleration only if 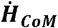 is constant. However, a CoP shift is likely to cause a change in angular momentum, as follows from the equation of rotational motion of a system of linked rigid segments about its CoM:

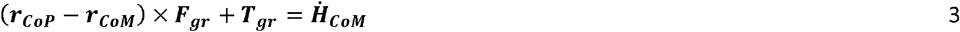

The above triggers the question why analyses using the vector between ***CoP*** and ***CoM’*** have been successful in making predictions about control of the CoM in standing and walking. An explanation for this is suggested by combining equations 2 and 3. Equation 2 implies that a change in linear momentum requires a change in ***F***_***gr***_, which has, in general, also consequences for the angular momentum (equation 3). If angular momentum is controlled to be close to zero during walking, as has been suggested (Herr and Popovic, 2008), this implies that is close to zero, which simplifies equation 3 to. equals 0 around the sagittal and transverse axes. Thus, the horizontal ground reaction forces are closely approximated by linear functions of the horizontal distances between CoM and CoP in the corresponding directions (for explanation see supplementary material).

In walking, ***H***_***CoM***_ is not exactly zero; it varies quasi-periodically (Bennett et al., 2010; Negishi and Ogihara, 2023). However, if horizontal linear momentum in a given plane is cross-correlated with angular momentum around the axis perpendicular to that plane, the equations of motion still predict a strong correlation between horizontal ground reaction force and the distance between CoM and CoP in the same plane (supplementary material). For brevity, we will henceforth refer to an angular momentum around the transverse axis as an angular momentum in the sagittal plane and to an angular momentum around the sagittal axis as an angular momentum in the frontal plane, as we will associate these to linear momenta in the same planes.

We hypothesized that, in human walking, linear and angular momentum in the same plane follow correlated quasi-periodic functions, such that also horizontal distances from CoM to CoP and horizontal ground reaction forces are correlated. Given the destabilizing effect of gravity, maintaining quasi-periodic patterns of linear and angular momentum requires control of both momenta through either passive mechanics or neuromuscular feedback. Therefore, we also hypothesized that deviations in linear and angular momentum from their planned trajectory (which we assume to be the average pattern over gait cycles) are negatively correlated with respectively deviations in horizontal ground reaction forces and deviations in ground reaction force moments around the CoM later in the gait cycle as follows:

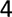

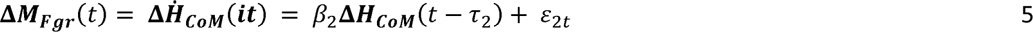

with *ε* representing time-dependent errors and ***M***_***Fgr***_ **= (*r***_***CoP***_. **− *r***_***CoM***_) × ***F***_***gr***_.

We used existing data 1) to assess the temporal patterns of linear and angular momentum and their correlation, 2) to assess the correlation between the distance from CoM to CoP and the horizontal ground reaction force in the corresponding direction, and 3) to verify the existence of negative correlations as described by equations 4 and 5. These analyses show why using the distance from CoM to CoP does provide insight into control of linear momentum even though the relationship between this distance and horizontal ground reaction forces is not a given mechanically. Moreover, they illustrate that linear and angular momentum are both controlled during gait and the control of the one constrains control of the other.

## Methods

### EXPERIMENTAL DATA

Ground reaction forces and whole-body CoM trajectories during treadmill walking were obtained from healthy young adults, participating in a previous experiment (van Leeuwen et al., 2020). Data from 13 participants (10 females, 3 males; mean (SD) age: 30 (9) yrs, mass 73 (17) kg, length 1.71 (0.09) m), for whom complete data for normal and slow walking speeds were available, were included in the analysis. In short, participants walked on an instrumented dual-belt treadmill (Motek-Forcelink, Amsterdam, Netherlands) at normal and slow (1.25 and 0.6125 × √(leg length) m/s) normalized (Hof, 1996) speeds. Stride frequency was imposed by means of a metronome, at the average frequency during the final 100 steps of a familiarization trial. Trials lasted five minutes for normal walking and ten minutes for slow walking. Ground reaction forces were recorded from force plates embedded in the treadmill and sampled at 200 Hz. Full body kinematics were measured using Optotrak (Northern Digital Inc, Waterloo Ontario, Canada) sampling at 50 samples/s. Cluster markers were attached to the feet, shanks, thighs, pelvis, trunk, upper arms and forearms. Corresponding anatomical landmarks were digitized using a six-marker probe.

### DATA ANALYSIS

For all trials, we analyzed 200 consecutive strides. Heel strikes were detected based on the CoP derived from force plate data (Roerdink et al., 2008). Mass and moment of inertia of each segment were estimated using segment length and circumference as predictors and regression coefficients based on sex (Zatsiorsky et al., 1990). All calculations were done in a stationary, lab-based coordinate system. The segment’s center of mass location was estimated to be at a percentage of its longitudinal axis (de Leva, 1996; Zatsiorsky et al., 1990). The CoM was estimated as the weighted sum of the centers of mass of the segments and its numerical derivative was used as an estimate of CoM velocity. This was multiplied by body mass to obtain linear momentum. Whole-body angular momentum was calculated as the sum of segmental angular momenta relative to the whole-body CoM. Subsequently, we calculated the moment of the ground reaction force around the CoM as the time-derivate of the whole-body angular momentum as an estimate of, since calculation based on the ground reaction is highly sensitive to errors in relative positions of CoP and CoM.

For our first aim, we additionally calculated the correlation and cross-correlation between the time series of the linear and angular momentum in each plane.

For our second aim, we calculated coefficients of correlation between the time series of the distance from CoM’ to CoP and of the horizontal ground reaction forces for the mediolateral and anteroposterior directions separately.

For our third aim, to assess whether deviations in linear and angular momentum negatively predict future deviations in horizontal ground reaction forces and moments of the ground reaction force around the CoM, we first high-pass filtered all data at 0.1 Hz, to discard low frequency fluctuations in gait speed, apparent as drift of the position of the person on the treadmill. We considered the target values of ***p***_***com***_ and ***H***_***com***_ to have quasi-periodic patterns, which we estimated from the average time-normalized patterns over all gait cycles in a trial. Next, we predicted deviations in linear and angular momentum from the averaged patterns by fitting the models described by equations 4 and 5, with time t replaced by phase of the gait cycle i. using linear regression. Both models were fit in a least-squares sense to the n observations (time-normalized strides) at each phase i. Models were fit separately for the frontal and sagittal plane. Assuming phase-dependent feedback (e.g., Berger et al., 1984; Bruijn et al., 2015; Golyski et al., 2021; Magnani et al., 2021), we assessed the models separately for every phase *i* ≥ 50%. We started at 50% of the stride to allow for using data during the preceding half of the gait cycle. We normalized strides based on left heel strikes. Normalizing strides from right heel strike to right heel strike provides complementary, but similar information and is hence not reported here. The value for the delay could approach zero for corrections based on intrinsic mechanics or amount to several hundreds of ms for active corrections that take effect for example after foot placement. These delays may vary between individuals and gait speeds. Thus, for each trial we optimized the delay across all phases in the gait cycles of a trial. We fitted the models for all possible delays from 1 to 49% of the gait cycle and retrieved the model that yielded the largest average negative correlation. For every resulting model we checked for which phases of the gait cycle the model fit was significant.

## Results

Linear and angular momentum in both the frontal plane and the sagittal plane showed cyclic patterns, with a similar frequency and phase (Figure 2). In the frontal plane, linear and angular momentum were highly correlated. In the sagittal plane, the momenta had a clear phase difference, resulting in low correlations but high cross-correlations (Figure 2).

**Figure 1.**
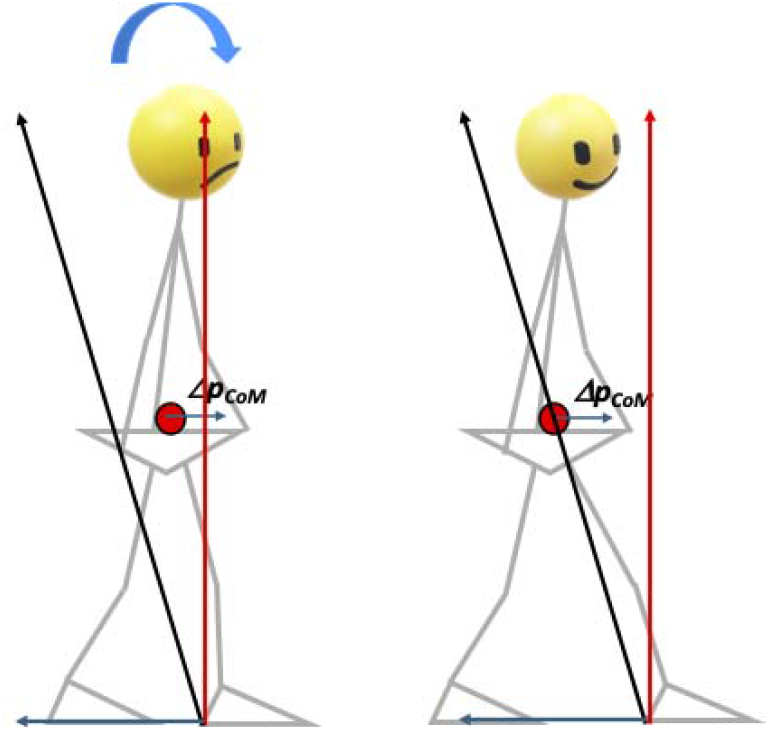
Illustration of the necessity to adjust the CoP if both linear and angular momentum have to be controlled. In both panels, the solid red circles represent the CoM which has a too high forward linear momentum **(Δp**_**CoM**_). To correct this momentum, a horizontal ground reaction force is needed (blue arrows). This causes an increase in forward angular momentum. At the same time the vertical ground reaction force (red arrows) causes a backward change in angular momentum. With the leading foot and consequently the CoP positioned as in the left panel, the resultant ground reaction force causes an increase in forward angular momentum (curved arrow). In the right panel, moments of vertical and horizontal ground reaction forces cancel, or in other words the resultant ground reaction force causes no moment around the CoM, keeping angular momentum constant.

**Figure 2.**
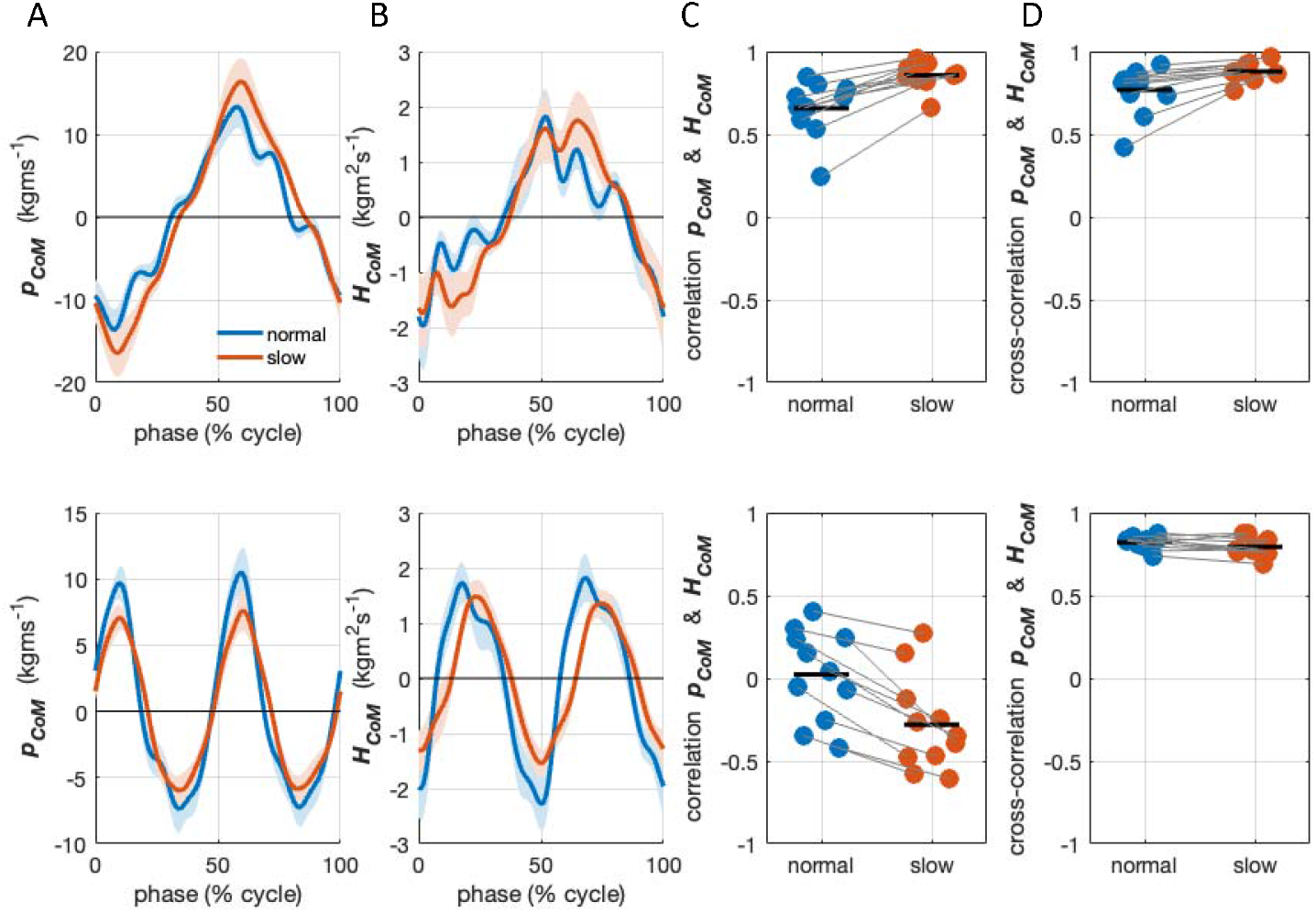
Momentum time series in normal and slow walking averaged over strides and participants (**A**) linear momentum and (**B**) angular momentum. (**C**) Coefficients of correlation between the time series of linear and angular momenta and (**D**) maximum cross-correlation coefficients between the time series of linear and angular momenta. Top panels represent frontal plane momenta and bottom panels represent sagittal plane momenta. The shaded areas indicate 95% confidence intervals. Each dot represents a single participant, the bars represent the mean values.

The distance between ***CoM’*** and ***CoP*** was strongly correlated to the horizontal ground reaction force for all participants, at both slow and normal speeds and in both directions (Figure 3).

**Figure 3.**
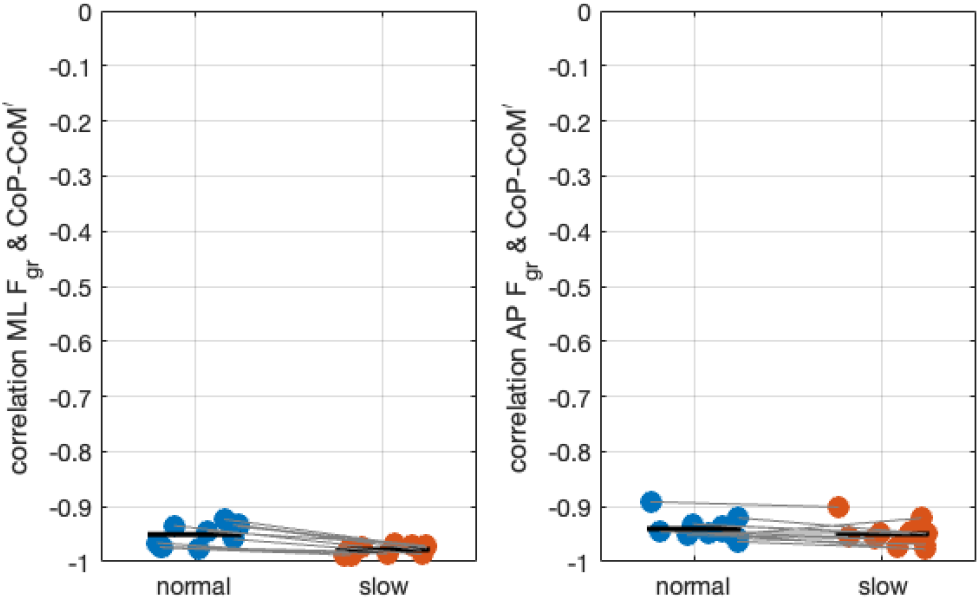
Coefficients of correlation between the horizontal ground reaction forces and distances from CoM’ to CoP in the frontal (left) and sagittal (right) plane, in normal and slow walking. Each dot represents a single participant, the bars represent the mean values.

In the frontal plane, deviations in ***p***_***com***_ and ***H***_***com***_ predicted deviations in 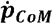 and 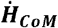 respectively, with significant negative regression coefficients throughout most of the gait cycle in normal and slow walking (Figure 4).

**Figure 4.**
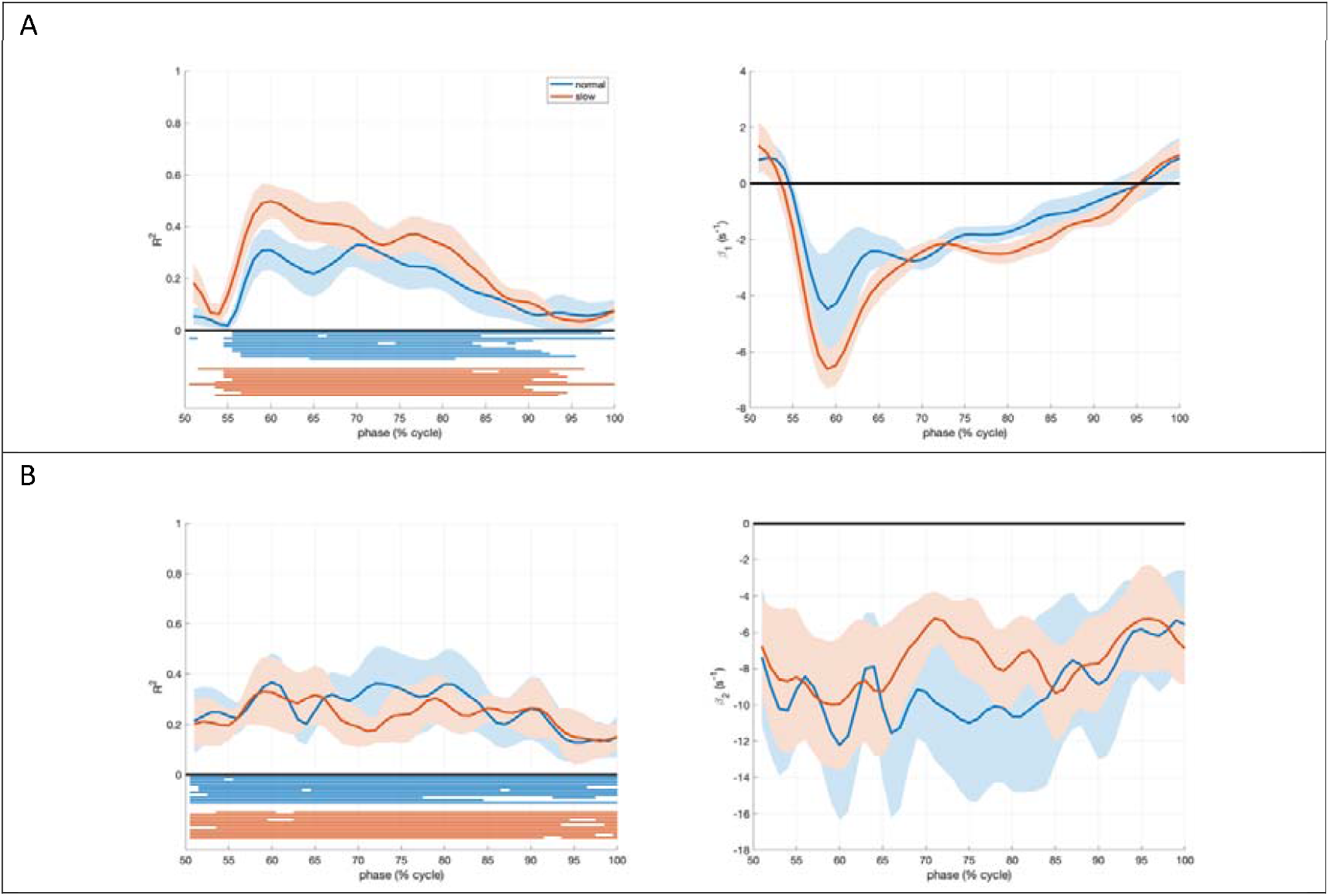
Models predicting deviations in the rate of change of linear and angular momentum in the frontal plane. **(A):** Model described in equation 4, relating deviations in mediolateral ground reaction force at phase *i* in the second half of a stride to deviations in the mediolateral impulse at a phase lead *τ*_1_ in normal and slow walking. The lines represent the averaged (over participants) phase-dependent goodness of fit (R^2^) and feedback gain (*β*_*1*_),, the shaded areas indicate the 95% confidence intervals. The horizontal lines in the left panels indicate for each individual participant and each speed when the model fit was significant with negative gains. (**B**) Model described in equation 5, relating deviations in 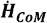 at phase *i* in the second half of a stride to deviations ***H***_***CoM***_ in the sagittal plane at a phase lead *τ*_2_ in normal and slow walking. The lines represent the averaged (over participants) phase-dependent goodness of fit (R^2^) and feedback gain (*β*_2_), the shaded areas indicate the 95% confidence intervals.

Also in the sagittal plane, deviations in ***p***_***com***_ and ***H***_***com***_ predicted deviations in 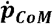 and 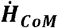 respectively, with significant regression coefficients throughout most of the gait cycle (Figure 5).

**Figure 5.**
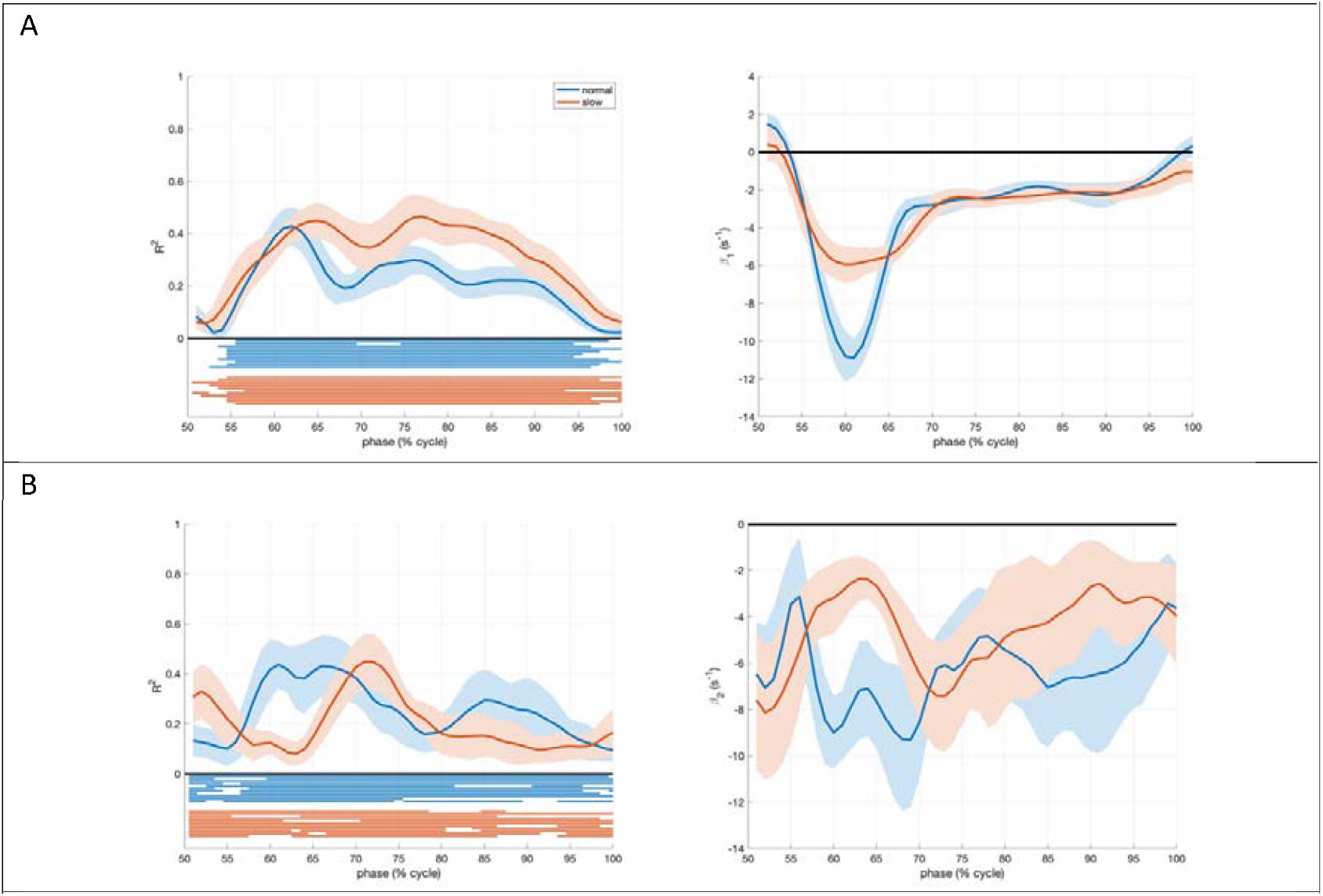
Models predicting deviations on the rate of changes of linear and angular momentum in the sagittal plane. (**A**): Model described in equation 4, relating deviations in mediolateral ground reaction force at phase *i* in the second half of a stride to deviations in the mediolateral impulse at a phase lead *τ* _1_ in normal and slow walking. The lines represent the averaged (over participants) phase-dependent goodness of fit (R^2^) and feedback gain (*β*_*1*_), the shaded areas indicate the 95% confidence intervals. The horizontal lines in the left panels indicate for each individual participant and each speed when the model fit was significant with negative gains. (**B**) Model described in equation 5, relating deviations in 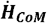 at phase *i* in the second half of a stride to deviations ***H***_***CoM***_ in the sagittal plane at a phase lead *τ*_2_ in normal and slow walking. The lines represent the averaged (over participants) phase-dependent goodness of fit (R^2^) and feedback gain (*β*_2_), the shaded areas indicate the 95% confidence intervals.

## Discussion

As mentioned in the introduction, the horizontal location of the CoP relative to the CoM (***r***_***CoP***_***-r***_***CoM’***_) and related concepts like foot placement have been interpreted as reflecting actions to control whole-body linear momentum. We here argued that mechanically speaking this is incorrect. Apart from Hof’s suggestion based on equation 1, the use of ***r***_***CoP***_***-r***_***CoM***_ appears to have been derived from models representing the human body by a mathematical inverted pendulum, see for example for standing (Morasso and Schieppati, 1999) and for walking (Townsend, 1985). A mathematical pendulum does not have angular momentum around its CoM, hence the ground reaction force points at the CoM, or in other words the external moment is always zero. The human body has a distributed mass, and hence it also has a time-varying angular momentum in standing and walking. As such, for adequate control it is necessary (but not sufficient) that both linear and angular momentum are controlled. In spite of the erroneous or over-simplified mechanical rationale underlying the use of ***r***_***CoP***_***-r***_***CoM***_, the distance between CoM’ and CoP has successfully been used to predict linear CoM acceleration from experimental data of human standing (Morasso et al., 1999; Winter, 1995). Here, we showed that also in experimental data from human walking the distance between CoM’ and CoP is strongly predictive of the horizontal ground reaction force (Figure 3).

A strong correlation between horizontal CoM-CoP distance and ground reaction force was expected if whole-body linear and angular momentum would be cross-correlated with small phase differences. Both expectations were confirmed by our analyses, and this may explain the success of using the CoM-CoP distance and related concepts, like foot placement. In addition, it is likely that constraints arising from the leg’s anatomy, force transmission between the foot and the surface, and the presence of a large vertical force component (on average equalling body weight) contribute to the correlation between horizontal CoM-CoP distance and ground reaction force. These constraints imply that the variance in direction of the forces applied to the ground by a foot in a particular orientation of the lower limb relative to the rest of the body is limited. For example, when standing upright, it is impossible to exert a substantial medio-lateral force on the ground. When a substantial medial ground reaction force is required, one must first move the foot laterally, resulting in a shift in CoP.

Regression equations 4 and 5 do not necessarily reflect neural feedback. Passive mechanics may contribute to coupling of the ground reaction force and the external moment to the preceding state of the body (Patil et al., 2019) and will have a delay that does not reflect the time needed for neural transmission. In fact, this can even occur without delay. As can be seen in figures 4 and 5 R^2^-values averaged between 0.2 and 0.4, indicating 20-40% of variance explained. We did not expect perfect correlations, in view of effects of measurement errors and neuromuscular noise on both the independent and dependent variables in the models. In addition, rates of change in momenta, the dependent variables are in part caused by gravity. The fact that significant negative correlations were found in all participants is in our view strong support for simultaneous control of both linear and angular momentum. We have previously proposed that for linear momentum a similar regression approach can be used to identify the lumped effect of passive and active control (van Dieën et al., 2024). In this case, CoM position and velocity were combined to predict horizontal ground reaction forces. We used a simpler approach here, to allow direct comparison between the control of linear and angular momentum. We suggest that the models presented here can be optimized to identify the control of linear and angular momentum. This would involve optimizing the number of state variables included as predictors, the estimation of the delay, and estimation of segmental and whole-body CoM locations, as well as correcting for effects of gravity on ground reaction forces and moments (Geng et al., 2024).

Given the many degrees of freedom of the human musculoskeletal system, it is not clear how simultaneous control of angular and linear momentum is realized. Multiple solutions allow for this, but not all of these will be desirable, from for example an energetic perspective, or even in keeping with the aim of locomotion. Ultimately, momenta can only be controlled through forces and moments acting on body segments. The first term of equation 1 has been related to the effects of ankle moments and foot placement as both affect ***r***_***CoP***_ and ***F***_***gr***_ (Hof, 2007; van Leeuwen et al., 2022). However, controlling both ***r***_***CoP***_ and ***F***_***gr***_ such that 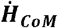 is also controlled cannot be achieved by a single joint moment, as this would leave one moment to control two outputs in each plane. Similarly, foot placement can be considered as a single control variable for each plane and is thus insufficient to control two outputs. As illustrated in the results section, when a change in linear momentum is needed and hence a change in ***F***_***gr***_, a simultaneous change of the CoP is needed to avoid an undesired change in ***H***_***com***_. Of course, this argument can be reversed. Shifting the CoP relative to the CoM allows decoupling of the control of linear and angular momentum. The regression coefficients for the association between linear momentum and subsequent horizontal forces (Figure 4a and 5a) indicated strong corrections of linear momentum in early stance. This may reflect that linear momentum can be corrected in this phase of the gait cycle without undue consequence for angular momentum by choosing an appropriate foot placement location. But, also passive mechanics, specifically effects of collision forces can play a role. Effects of collision forces, mechanically determined in part by CoM states right before the collision, would be manifested in early stance. This may be especially relevant for foot placement in the sagittal plane, as collision forces can provide stability to passive dynamic walkers in the sagittal plane (Mcgeer, 1990) but not in the frontal plane (Bauby and Kuo, 2000). Maximum gains for linear momentum control were found in this phase of the gait cycle. Moreover, position gains in the sagittal plane appeared to be speed dependent, which could be accounted for by differences in collision forces between speeds.

This study was limited to steady-state walking, affected only by minor perturbations such as neuromuscular noise. It is safe to say that both linear and angular momentum must be controlled during other gaits such as running, stair negotiation and perturbed or non-steady-state gait, but how these momenta vary over time remains to be shown. During gait initiation specifically, linear and angular momentum are not correlated (Bechet et al., 2025). This implies that the correlation between CoP-CoM distance and horizontal force does not necessarily hold during this task. Furthermore, this task may be less under feedback control than steady-state gait (Tisserand et al., 2018).

We showed that corrections of linear momentum are constrained by angular momentum. Obviously, the reverse also holds, corrections of angular momentum may affect linear momentum. Experimental data on perturbed gait show that control of angular momentum to remain upright may require linear momentum to be temporarily ‘sacrificed’, i.e. brought further from the unperturbed state. Specifically, after a trip, which induces a substantial change in angular momentum, recovery responses caused a further perturbation of forward linear momentum, but restored angular momentum (Pijnappels et al., 2004). Conversely in standing on a narrow beam, where shifts of the CoP are constrained, linear momentum is controlled through hip moments, allowing changes in angular momentum (Otten, 1999). We conclude that studying the control of linear momentum and angular momentum separately has limitations, because linear and angular momentum are mechanically coupled as both depend on external forces (like ***F***_***gr***_), while in addition anatomical constraints and foot-ground interactions generally require adjusting foot placement for the control of linear and angular momentum.

## Nomenclature

All kinematic variables are expressed with respect to an inertial reference frame affixed to the world. All moments of inertia are with respect to the center of mass of the respective segment/body. Dot notations denote derivatives with respect to time.
a_CoM’_ = linear acceleration of the projection of body center of mass on the support surface. CoM = whole-body center of mass.
CoM’ = projection of the whole-body center of mass on the support surface.

CoP: center of pressure, or point of application of the ground reaction force.
F_gr_: ground reaction force vector.
g: gravitational constant.
H_CoM_: angular momentum of the whole body around its center of mass.
I_i_: inertia tensor of segment i.
m: body mass.
m_i_: mass of segment i.
T_gr_: torque around the vertical axis due to a couple of horizontal ground reaction forces
p_CoM_: linear momentum of the whole body.
r_CoM_: position vector of the body center of mass.
r_CoM’_: position vector of the projection of the body center of mass on the support surface.
r_CoP_: position vector of the point of application of the ground reaction force or center of pressure.
r_i_: position vector of the center of mass of segment i relative to the body center of mass.
v_CoM_: velocity of the body center of mass.
v_i_: velocity of segment i relative to the body center of mass.
ω_i_: angular velocity of segment i.

## Supplementary material

Under the assumption that ***H***_***CoM***_ is close to zero throughout the gait, which implies that also 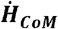 should be close to zero, we can readily predict the horizontal ground reaction force. Using a right-handed coordinate system, where positive *x* is forward, positive *y* is upward, and positive *z* is right, we define(***r***_***CoP***_ − ***r***_***CoM***_)= [*x y z*]’. Consider a correction of linear momentum of the CoM in the fore-aft direction (*F*_*x*_) and no correction in the mediolateral direction (*F*_*z*_ = 0)., Furthermore, we assume *y* and thus *F*_*y*_ to be constant (mg). Thus ***F***_***gr***_ = [*F*_*x*_ *mg 0*]’. If the change in angular momentum about the z-axis is constrained to be zero, the equation of rotational motion simplifies to:

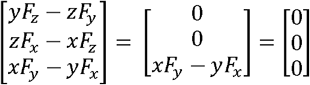

This shows that, the horizontal ground reaction force is proportional to *x*, the distance between CoM’ and CoP. Obviously, any deviation from 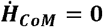, or from the assumption that the vertical ground reaction force is constant, will cause an error in the prediction of the horizontal ground reaction force from the distance between CoM’ and CoP.

Variance of the vertical ground reaction force during walking is limited. Horizontal and angular momentum in each plane show cyclic patterns with a similar frequency and phase (see Figure 2 in the main text). Therefore, so do their derivatives: the horizontal ground reaction forces and the moments of the ground reaction force. Based on these findings, we ran simulations of the equations of linear and rotational motion. For these simulations, 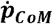 and 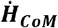 were defined as 1Hz sinusoids with amplitudes of 40 N and 14Nm respectively (resembling the experimental data). *F*_*x*_ was calculated from the equation of linear motion, *F*_*y*_ was considered constant and mass was set to 70 kg, *F*_*z*_ was set to zero. ***F***_***gr***_ was inserted into the equation of rotational motion with 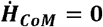, to obtain CoP-CoM’.

The simulations yielded negative linear relationships between ***r***_***CoP***_***-r***_***CoM’***_ and the horizontal component of ***F***_***gr***_ (Figure S1). With limited differences in phase between linear and angular momentum (as demonstrated in the main text), the correlation remains close to −1.

**Figure S1.**
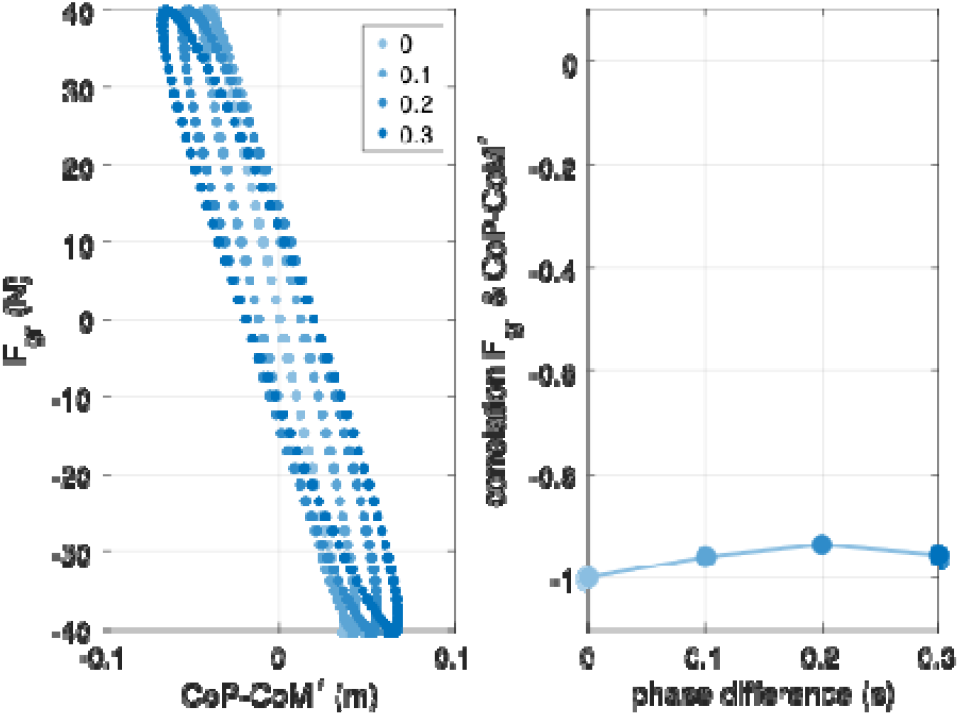
Results of simulations of the equations of linear and rotational motion. 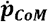 and 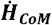 were defined as 1Hz sinusoids with amplitudes of 40 N and 14Nm respectively (resembling experimental gait data). ***F***_***x***_ was calculated from the equation of linear motion, ***F***_***y***_ was considered constant and mass was set to 70 kg, ***F***_***z***_ was set to zero. ***F***_***gr***_ was inserted into the equation of rotational motion to obtain CoP-CoM’. The results, show the strong negative correlation between ***r***_***CoP***_***-r***_***CoM’***_ and ***F***_***gr***_ and its limited sensitivity to differences in phase (ranging from 0 to 0.3 s) between 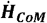 and ***F***_***gr***_.

